# Mouth Function Determines The Shape Oscillation Pattern In Regenerating *Hydra* Tissue Spheres

**DOI:** 10.1101/565465

**Authors:** R. Wang, T. Goel, K. Khazoyan, Z. Sabry, H.J. Quan, P.H. Diamond, E.M.S. Collins

## Abstract

*Hydra* is a small freshwater polyp capable of regeneration from small tissue pieces and from aggregates of cells. During regeneration, a hollow bilayered sphere is formed that undergoes osmotically driven shape oscillations of inflation and rupture. These oscillations are necessary for successful regeneration. Eventually, the oscillating sphere breaks rotational symmetry along the future head-foot axis of the animal. Notably, the shape oscillations show an abrupt shift from large amplitude, long period oscillations to small amplitude, short period oscillations. It has been widely accepted that this shift in oscillation pattern is linked to symmetry breaking and axis formation. However, recent work showed that regenerating tissue pieces inherit the parent animal’s body axis and thus are asymmetric from the beginning. Thus, there is no mechanistic explanation for the observed shift in oscillation pattern and no clear understanding of its significance for *Hydra* regeneration. Using *in vivo* manipulation and imaging, we quantified the shape oscillation dynamics and dissected the timing and triggers of the pattern shift. Our experiments demonstrate that the shift in the shape oscillation pattern in regenerating *Hydra* tissue pieces is caused by the formation of a functional mouth, thereby linking morphological readouts to physiologically relevant events during regeneration. This study shows the power of using modern experimental techniques to revisit old questions in pattern formation and development.

## INTRODUCTION

*Hydra* is a small (~1 cm long), transparent, radially-symmetric freshwater cnidarian polyp (Fig 1A). It consists of a cylindrical body column with a tentacle ring and a dome-shaped hypostome containing the mouth on one end and a foot that anchors the animal to the substrate on the other. *Hydra* is composed of only two tissue layers, an outer ectodermal epithelium and an inner endodermal epithelium, separated by a basal lamina called the mesoglea. Body shape is regulated by contractile processes on the epithelial cells called myonemes, which are oriented longitudinally along the head-foot axis in the ectoderm and circumferentially in the endoderm (1). This simple anatomy, combined with the ability to regenerate a complete polyp from tissue pieces and from aggregates of body column cells, made *Hydra* an important model system for biologists and physicists alike to study regeneration, axis formation, and patterning (2).

**Figure 1.**
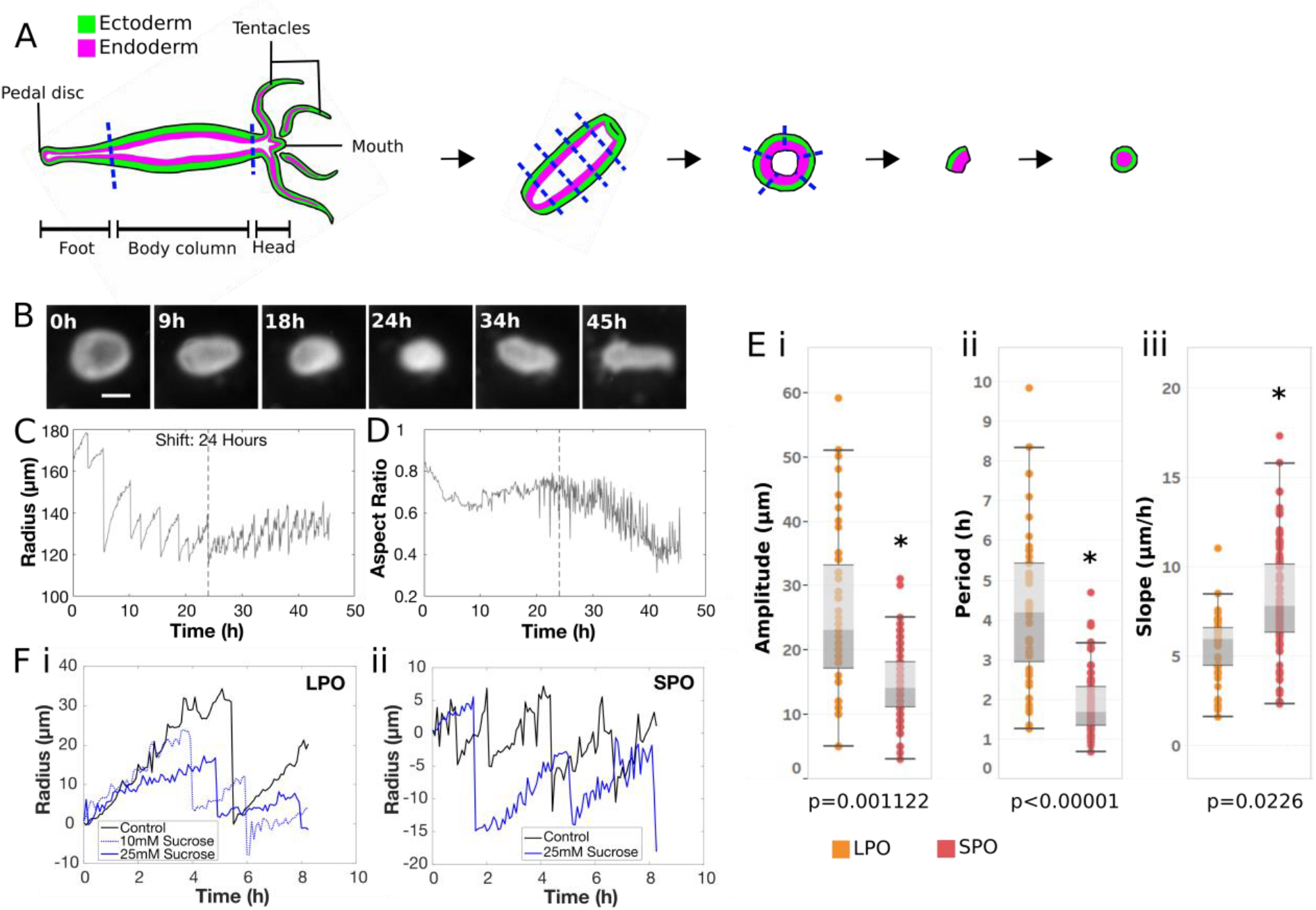
Generation of tissue spheres and quantification of oscillation dynamics. A. Preparation of tissue pieces from a *Hydra* polyp (see Materials and Methods). B. Representative images of regenerating tissue spheres at various time points during regeneration. In the 45 h image, the regenerated head with tentacles is to the left. Scale bar 150 µm. C. Plot of effective radius, calculated as the radius of a circle with an area equal to that of the tissue piece, as a function of time for the sphere shown in B. D. Plot of aspect ratio as a function of time for the same tissue sphere. Dashed line indicates the time of shift from LPO to SPO. E. Box-whisker plots of i. amplitudes, ii. time periods and iii. slopes for LPOs and SPOs. Asterisks indicate a statistically significant difference from LPO (p<0.05): amplitude p = 0.00112, period p < 1e-5, slope p =0.0226. F. Plot of effective radius as a function of time at different sucrose concentrations in the external medium during i. LPOs and ii. SPOs.

One of the earliest attempts at modelling axial patterning in *Hydra* was made by Alfred Gierer and Hans Meinhardt, who proposed a reaction-diffusion model consisting of a short range head activator, a long range head inhibitor, and a gradient for the activator source (3). The model qualitatively explains pattern formation from a homogeneous starting state. However, a lack of quantitative experimental data has limited progress on validation and refinement of this and subsequent models (4). Recently, the availability of a fully sequenced genome (5), various transgenic reporter lines (6, 7) and CRISPR genome editing tools (8) has allowed researchers to reexamine earlier models and studies of *Hydra* regeneration and gain new insights.

Here, we revisit a striking phenomenon that occurs during *Hydra* regeneration from tissue pieces (9) and aggregates of cells (3). As they regenerate, both tissue pieces and aggregates form a hollow bilayered sphere with ectodermal cells on the outside and endodermal cells on the inside. These *Hydra* spheres undergo osmotically driven cycles of swelling and subsequent rupture, referred to as shape oscillations (10). Shape oscillations are sawtooth shaped, consisting of cycles of long inflation phases, followed by an abrupt deflation of the sphere due to local tissue rupture (11). The inflation phase is caused by the uptake of water and the active pumping of sodium ions into the lumen of the sphere (12). Initially, inflation is isotropic. The *Hydra* sphere’s aspect ratio, defined as the ratio of the short axis to the long axis of an ellipse fit to the sphere, is close to unity. As time progresses, the swelling becomes increasingly anisotropic - the aspect ratio decreases with sharp dips during deflation of the hollow sphere. The regenerating animal establishes a body axis and develops a mouth and tentacles by approximately 48 hours (13). Critically, the oscillations have been reported to show an abrupt shift in frequency that coincides with shape symmetry breaking, defined as a decrease in aspect ratio.

To the best of our knowledge, the mechanism underlying this shift in oscillation pattern has not been determined. It has long been hypothesized, however, that the oscillation pattern shift is linked to both shape symmetry breaking, and the morphogen-driven patterning and axis determination proposed by Gierer and Meinhardt. Sato-Maeda and Tashiro were the first to probe this connection two decades ago. They reported the sawtooth shape of the oscillations, and described a method of detecting shape symmetry breaking in cell aggregates by quantifying the divergence of orthogonal radii as the regenerating animal elongated along one axis (11). This approach represented a measure of body axis formation that could be quantitatively linked to other morphological fluctuations. Fütterer *et al.* subsequently analyzed the shape of regenerating *Hydra* spheres originating from tissue pieces in greater detail, using Fourier decomposition to reveal 3 distinct temporal stages: 1. Large amplitude long period oscillations (LPO) of the zeroth mode (size of the tissue piece), 2. Small amplitude, short period oscillations (SPO) of the zeroth mode associated with fluctuations of the second mode (elongation), 3. Strong increase in the second mode during contractions. They reported that shape anisotropy always occurred after the completion of LPOs, suggesting a correlation between oscillation dynamics and formation of the body axis (13).

*Hydra* spheres derived from cell aggregates and from small tissue pieces exhibit similar oscillation dynamics. It was also reported that regenerating spheres developed their body axes in alignment with an applied temperature gradient regardless of their origin (14). Consequently, it was conjectured that both tissue pieces and cell aggregates begin from a homogenous state and must break symmetry *de novo*. This idea was supported by the finding that the time of pattern shift from LPO to SPO coincides with the emergence of criticality in the patch size distribution of the *Hydra* head-specific gene *ks1* (14). Furthermore, the timing of oscillation pattern shift was comparable to the timing of the emergence of Wnt3 expression patches in larger cell aggregates, at approximately 24 h (15). Because Wnt3 is the earliest known marker expressed during head regeneration in *Hydra*, (16, 17) Soriano *et al.* concluded from these findings that the oscillation pattern shift coincides with the establishment of biochemical asymmetry. Thus, the pattern shift was regarded as a reliable and easily detectable morphological marker for symmetry breaking and axis specification in both aggregates and small tissue pieces (14).

Because of this apparent link, subsequent theoretical models by Soriano *et al.* (18) and Mercker *et al.* (4) coupled tissue mechanics with reaction-diffusion of morphogens to explain axis formation in *Hydra* and equated the time of oscillation pattern shift with the time of symmetry breaking and body axis specification to constrain their models. However, it was recently shown that spheres derived from small tissue pieces, unlike aggregates, inherit the parental body axis in the form of myoneme organization (19). Therefore, spheres derived from tissue pieces have a prespecified body axis and the observed oscillation pattern shift cannot be a marker of an axis formation event. Thus the underlying cause and relevance of the shift from LPOs to SPOs remains to be determined.

Here, we use *in vivo* manipulation and imaging to quantify shape oscillation dynamics and experimentally dissect the timing and triggers of the pattern shift. First, we demonstrate that both LPOs and SPOs are driven by osmotic pressure, suggesting that the observed differences do not arise from different swelling mechanisms but from changes in the local yield strength of the tissue spheres. Consistent with this idea, we find that the site of tissue rupture is random during LPOs, whereas it is conserved during SPOs, suggesting the existence of a fixed mechanical weak point during SPOs. Because regenerating tissue pieces derived from nerve-free animals, which are unable to open their mouths, exhibit only LPOs, we conclude that it is not the existence of a mouth structure but rather mouth function that gives rise to the shift in oscillation pattern.

Indeed, tissue pieces with an intact, functional mouth were found to exhibit only SPOs, whereas tissue pieces from the heads of nerve-free animals with a structurally normal mouth only exhibit LPOs. Together, these experiments demonstrate that the shift in oscillation pattern observed in regenerating *Hydra* tissue pieces indicates the onset of mouth function, providing an easily observable readout for an important regeneration milestone. In addition to providing a mechanistic explanation for shape oscillation dynamics, this study also allowed us to estimate a lower bound for the tissue yield strength, a parameter which may prove useful for future models of *Hydra* regeneration.

## MATERIALS AND METHODS

### Hydra strains and culture

*Hydra vulgaris* strain AEP, *Hydra vulgaris* (formerly *Hydra magnipapillata*) strain sf-1 (temperature sensitive interstitial stem cells), *Hydra vulgaris* strain A10 (chimera consisting of *Hydra vulgaris* (formerly *Hydra magnipapillata* strain 105) epithelial cells and sf-1 interstitial cells) (20) and *Hydra vulgaris* “watermelon” (AEP expressing GFP in the ectoderm and DsRed2 in the endoderm) (7) were used for experiments. Polyps were kept in *Hydra* medium (HM) composed of 1 mM CaCl_2_ (Spectrum Chemical, New Brunswick, NJ), 0.1 mM MgCl_2_ (Sigma-Aldrich, St. Louis, MO), 0.03 mM KNO_3_ (Fisher Scientific, Waltham, MA), 0.5 mM NaHCO_3_ (Fisher Scientific), and 0.08 mM MgSO_4_ (Fisher Scientific) prepared with MilliQ water, with pH between 7 and 7.3, at 18°C in a Panasonic incubator (Panasonic MIR-554) in the dark. The *Hydra* were fed three times per week with *Artemia* nauplii (Brine Shrimp Direct, Ogden, UT). Animals were cleaned daily using published procedures (21).

### Generation of nerve-free *Hydra*

Nerve-free *Hydra* were generated using either of two methods. Watermelon animals were made nerve-free as described by Tran *et al*. (22). Briefly, the animals were incubated in 0.4% colchicine (Acros Organics, Thermo-Fisher Scientific, Waltham, MA) in HM for 8 h in the dark. This 8 h incubation was then repeated 3 weeks following the first treatment. Colchicine-treated *Hydra* are susceptible to bacterial infection, so the animals were kept in HM supplemented with 50 μg/mL rifampicin (EMD Millipore, Burlington, MA) at 18°C in the dark in the incubator. Non-transgenic nerve-free animals were generated by heat shock treatment of the sf-1 and A10 strains (20, 23, 24). Sf-1 and A10 animals were heat shocked in an incubator at 29°C in the dark for 48 h and then moved back into the 18°C incubator. All nerve-free animals were force fed and “burped” as per the protocol described in Tran *et al.* (22).

### Preparation of tissue pieces

Tissue pieces were cut with a scalpel (Sklar Instruments, West Chester, PA) from the body columns of adult non-budding *Hydra* starved for 24 h, as shown in Figure 1A. The head was amputated immediately below the tentacles. A second cut was made above the foot to isolate the body column. Depending on the size of the resulting body column piece, one to three cross sectional cuts were made to extract rings. The rings were cut into 4 or more pieces and allowed to round up in HM for approximately 2 h – measured from the time of initial excision of the body column piece. Once rounded up, tissue pieces were selected by size (~200 µm diameter) for use in experiments (Fig. 1A).

### Preparation of head and foot tissue pieces

Head tissue pieces were prepared under a stereo microscope. The animal’s head was removed immediately below the tentacle ring, and then the tentacle bases were excised. The remaining head tissue pieces were given approximately 1 h to round up (measured from cutting the first tissue piece) and placed individually into custom made agarose wells for time-lapse imaging. Foot tissue pieces were prepared by cutting the animal immediately above the basal disc and allowing the resulting tissue to round up for 2 h. Subsequently, rounded pieces of the same approximate size as body column tissue pieces were selected and imaged.

### Imaging of shape oscillations

Regenerating tissue pieces were placed in agarose wells made using a 1% solution of agarose or low melting point agarose (Invitrogen, Carlsbad, CA) in HM. To make the wells, molten agarose solution was poured into 30 mm Falcon petri dishes (Thermo Fisher Scientific) and a comb with 1 mm wide teeth was placed vertically into the dishes to create wells. Once the agarose had solidified, the comb was removed, the wells were filled with HM and the tissue pieces were moved into the wells using a pipette. Imaging was accomplished using an Invitrogen EVOS FL Auto 2 microscope (Thermo Fisher Scientific) and the Invitrogen EVOS FL Auto 2.0 Imaging System software. Images were acquired every 5 minutes and stored as TIFF files. Viability of the tissue pieces was assayed by observing the presence of a body axis at 48 h, and the formation of tentacles and mouth opening upon presentation of *Artemia* at 96 h.

### Altering osmolarity of *Hydra* medium

To test the effect of changes in osmotic pressure on regenerating tissue pieces, tissue pieces were prepared and imaged as described above. However, the tissue pieces were kept in sucrose supplemented HM for imaging instead of HM. Sucrose (Sigma-Aldrich) was added to HM to final concentrations of 10 mM or 25 mM. Rifampicin (EMD Millipore) was added to a final concentration of 50 µg/mL to prevent bacterial growth in the presence of sucrose.

### Injections of microbeads and rupture site tracking

Tissue pieces were incubated at room temperature until at least 5 h after cutting to allow them to round up and form an internal cavity. An agarose trough for microinjection was cast as previously described (25). Hollow tissue spheres were placed in the trough in HM and injected with 1 µm green fluorescent (excitation/emission: 468/508 nm) microbeads (Thermo-Fisher G0100) using a WPI Pneumatic PicoPump - PV 820 (Sarasota, FL) and needles pulled using a Sutter Instrument P-1000 (Novato, CA). Successfully injected spheres were placed in agarose wells and imaged for 24 h as described above. The resulting videos were used to determine the location of rupture events by tracking the locations of ejection of fluorescent beads relative to a fixed feature on the sphere. The smaller of the two angles between the fixed feature and the rupture location was recorded.

### Visualization of myoneme arrangement in the head

Nerve-free *Hydra* prepared by heat shock treatment of strain A10 and untreated controls were fixed and stained with rhodamine-phalloidin (Biotium, Fremont, CA). The polyps were relaxed in 1 mL of 1 mM linalool (Sigma-Aldrich) in HM for 10 minutes and then fixed in 4% paraformaldehyde (Fisher) in HM for 20 minutes at room temperature. They were washed with HM thrice for 10 minutes each before being incubated overnight at 4°C in rhodamine-phalloidin diluted 1:100 in HM. The fixed stained samples were washed five times for 10 minutes each with HM. They were then placed on 22 mm × 40 mm glass coverslips (Corning Inc., Corning, NY) which had a piece of double-sided tape (3M, Maplewood, MN) running along the short edges of the coverslips. These coverslips were then sealed by placing 22 × 22 mm glass coverslips (Fisher Scientific) on top and the samples were imaged using an Olympus IX81 inverted microscope (Olympus Corporation, Tokyo, Japan) with an ORCA-ER camera (Hamamatsu Photonics, Hamamatsu, Japan). Slidebook version 5 (Intelligent Imaging Innovations, Denver, CO) was used to interface with the microscope and acquire z-stacks. Maximum intensity projections of the z-stacks were used to determine the orientation of the myonemes.

### Oscillation analysis

Images collected using the EVOS microscopes were opened in ImageJ (http://imagej.nih.gov/ij/, National Institutes of Health, Bethesda, MD, USA) and full regeneration was verified. Full regeneration was defined as the regenerated tissue piece exhibiting a well-defined body axis, head formation, and tentacle growth. Each image set was then analyzed using a custom Python script (Python 3.7.0, Python Software Foundation). The script first applies morphological image opening and closing to distinguish the sphere from the background, followed by watershed segmentation to detect and eliminate ejected cell debris. If the script failed to segment the raw image set, debris was removed from the images by manually tracing over the debris in ImageJ before analysis. For each image in a set, the script traces the boundary of the regenerating tissue to determine its area and effective radius and approximates its shape by fitting an ellipse and recording major and minor axes to determine the aspect ratio.

Subsequent processing and analysis of the data was carried out in MATLAB 2017b (MathWorks, Natick, MA, USA). The existence and timing of oscillation pattern shift in a data set was determined by having five researchers independently examine the radius-time plots for the data set and provide an estimate of the presence and timing of the shift. The data set was accepted as having a shift at a particular time if there was consensus of at least four of the researchers.

After shift presence and timing were determined, the amplitude, time period and slope of each oscillation was extracted. The amplitude was defined as the difference in maximum and minimum radius during the inflation phase. The time period was defined as the time difference between beginning of the inflation phase and end of the deflation phase. The swelling rate (slope) was obtained from a linear fit to the inflation phase of the oscillation. A two-sided Mann-Whitney U test was used to determine whether two sets of oscillation parameters originated from the same distribution. As individual oscillations within a single biological replicate cannot be considered independent, for each tissue piece the median values for each oscillation type were taken and used as inputs for testing. A p-value of 0.05 or lower rejects the null hypothesis that the two samples were drawn from the same distribution. For all conditions other than body column tissue pieces taken from wild-type animals regenerating in HM, the oscillations were classified as LPO or SPO based on comparison to wild-type LPO and SPO time periods.

Amplitudes and slopes were not used for this classification because these parameters can be affected by size and other properties of the tissue such as permeability and elasticity, which may vary depending on the source location (head, body columns, foot) of the sphere or treatments (nerve-free, sucrose) applied to it.

### Calculation of the yield strength of the tissue

The *Hydra* tissue sphere was treated as a linear elastic hollow spherical shell. Then, the elastic pressure experienced by the sphere is given by,

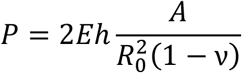

Here, E is the Young’s modulus of the tissue, ν is the Poisson’s ratio, h is the thickness of the shell, A is the amplitude of the sphere at the time of rupture and R_0_ is the minimum radius of the sphere. The tissue was assumed to be incompressible, so ν = 0.5. However, the results are not strongly dependent on the choice of ν. For example, if ν = 0.25 is used, as in Kücken *et al.* (10), the pressure is only reduced by a factor of 1.5. For the Young’s modulus, a value of 185 N/m^2^ was used, based on experiments by Veschgini *et al.*, (26) who measured the response of tissue spheres to uniaxial compression. The median values of minimum radius and amplitude were used, with R_0_ = 119 µm and A= 28 µm for LPOs and A=15.5 µm for SPOs, respectively. The shell thickness, h, was assumed to be 2 cell lengths. Cell size was estimated from measurements of cell area in maceration preparations (27) and from measurements of cell dimensions in histological sections (28). Approximating the cells as cubes, the former measurement was treated as the surface area of the cube and the latter was used to calculate the volume of the cube. From both measurements, cell size was estimated to be on the order of 10 µm. The thickness of the mesoglea, 0.3-3 µm, is an order of magnitude less compared to that of the cells (29, 30), and thus the mesoglea was neglected. The size of the hole caused by rupture was estimated from images that captured debris leaving the tissue sphere during a rupture event. The narrowest portion of the debris immediately adjacent to the sphere was averaged over three events and treated as an upper limit approximation of the size of the exit point, yielding a mean diameter of 26 µm, corresponding to 2-3 cell diameters.

### Comparison of the oscillation parameters to previously published values

Published histograms of the slopes during LPOs and SPOs and their mean values were taken from Soriano *et al.* (18). Because their distributions were non-normal with a left skew, the medians were smaller than the means. However, because the medians are difficult to extract from the plots without access to the data and the authors only reported mean values for some of the parameters, all parameter comparisons were performed on means and not medians. The mean values of time periods were obtained from visual inspection of the volume-*vs*-time plot in Soriano *et al.* (14). Average amplitudes were also obtained in the same manner from the radius-*vs*-time plot in Kücken *et al.* (10). As we only had access to published mean values, means were calculated for these parameters using our data to allow for better comparison. To calculate the means of our data, medians were first calculated for each biological replicate and then averaged over all the replicates.

## RESULTS AND DISCUSSION

As a freshwater animal, *Hydra* experiences a continuous inflow of water from the medium, through the tissues and into the gastric cavity (31, 32). The resulting internal pressure is periodically relieved by opening of the mouth (33). Regenerating *Hydra* spheres initially lack a mouth and therefore must relieve pressure from water accumulation by passive tissue rupture. This creates an oscillatory pattern of gradual osmotically driven swelling and rapid deflation due to tissue rupture and healing of the rupture site. These cycles of swelling and rupture show an abrupt shift in oscillation pattern from LPOs to SPOs, coincident with a change in the aspect ratio of the regenerating *Hydra* sphere.

### LPOs and SPOs have distinct oscillation parameters but a common driving mechanism

To examine the cause of the observed shift in oscillation pattern, we prepared tissue spheres (Fig. 1A) and imaged them over the course of regeneration. We only analyzed data from tissue spheres that regenerated fully, showing a defined body axis with head and tentacles (Fig. 1B). A shift in oscillation pattern was observed to coincide with a gradual decline in aspect ratio (Fig. 1C, D), as previously reported (10, 13, 18). From these radius-*vs*-time plots (Fig. 1C), we extracted amplitude, period, and swelling rate (slope) for LPOs and SPOs (see Materials and Methods) and found all parameters to differ significantly between the two oscillation types (Fig. 1E). A comparison of our data to the literature (10, 18) using the means of the oscillation parameters (see Materials and Methods) shows similar differences in these three parameters between LPO and SPO.

The oscillation amplitudes we observed correspond to cell shape changes of approximately 25% and 15% over the course of a single oscillation during LPOs and SPOs, respectively. While these are significant cell deformations, similar and more extreme deformations are observed during mouth opening in intact polyps over the course of tens of seconds (33). These numbers illustrate the remarkable deformability of *Hydra* tissue.

Previous studies and models assumed that both LPOs and SPOs are driven solely by osmotic pressure (4, 10). However, this was only experimentally tested for LPOs (10). To verify that SPOs are also osmotically driven, we incubated tissue pieces in hypertonic medium made by adding 10 mM or 25 mM sucrose to HM. Since the osmolarity between the inside and the outside of a tissue sphere equilibrates after several rupture events, we began incubation either 2 h post amputation to probe the effect of altered osmotic pressure on LPOs or 24 h post amputation to probe its effect on SPOs. Consistent with previous work (10), we observed a concentration-dependent decrease in swelling rates during the LPO cycle in the 2 h post amputation treatments (Table 1). Similarly, we obtained a negative correlation between slope and osmolarity in the 24 h post amputation treatment with 25mM sucrose. Moreover, the qualitative increase in slope from LPOs to SPOs was not affected by sucrose concentrations (Table 1), suggesting that the SPOs are also primarily osmotically driven, and that the increased rate of inflation is due to a secondary mechanism.

**Table 1.** Summary of oscillation parameters. Parameters are reported as the median of biological replicates with the first and third quartiles. * indicates significant difference from wild-type LPOs at p < 0.05. ** indicates significant difference from wild-type LPOs at p < 0.01. ^§^ indicates significant difference from wild-type SPOs at p < 0.05. ^§§^ indicates significant difference from wild-type SPOs at p < 0.01.

| | Period length (h) | Amplitude ($\mu\text{m}$) | Slope ($\mu\text{m/h}$) | Number of biological replicates |
| --- | --- | --- | --- | --- |
| **Wild-type (HM) LPO** | 4.2 (3.7, 5.8) §§ | 28.0 (21.0, 36.6) §§ | 6.3 (5.6, 7.1) § | 15 |
| **Wild-type (HM) SPO** | 1.8 (1.5, 1.9)** | 15.5 (13.0, 19.7)** | 7.5 (6.7, 9.8)* | 15 |
| **Sucrose 10mM 2h** | 2.3 (1.9, 3.0)** §§ | 13.5 (10.9, 19.1)** | 4.9 (4.4, 6.3) §§ | 14 |
| **Sucrose 25mM 2h** | 3.0 (2.8, 4.3) §§ | 14.3 (11.7, 19.8)** | 3.5 (3.1, 4.1)** §§ | 17 |
| **Sucrose 25mM 24h** | 4.1 (2.7, 4.5) §§ | 21.2 (13.5, 29.2) | 4.0 (3.6, 5.0)** §§ | 15 |
| **Wild-type head tissue piece** | 1.1 (0.9, 1.4)** §§ | 8.7 (6.3, 12.1)** § | 9.7 (7.4, 10.9)* | 4 |
| **Foot tissue piece** | 2.8 (2.3, 4.0) §§ | 21.0 (14.2, 30.3) | 7.6 (5.5, 7.9) | 5 |

The decrease in maximum amplitude of SPOs compared to LPOs indicates that the pressure required to trigger a rupture event has decreased. This can be explained either by the weakening of the tissue’s tearing strength (globally or locally), or the rupture becoming an actively controlled process. Both of these are attributes of the *Hydra* mouth. The mouth is a structural weak spot because it has a thinner mesoglea and an absence of myonemes running across it (34). The mouth also allows for active pressure release in the intact polyp, through the control of the nervous system (33). Soriano *et al.* proposed the first possibility (14), suggesting that the formation of a proto-mouth created a weak spot, but this idea was not tested experimentally.

### Rupture site becomes constant as regeneration progresses

To determine whether a fixed rupture site consistent with a permanent mechanical defect appears during regeneration, we used fluorescent microbead injection to visualize the rupture site in oscillating tissue spheres (Fig. 2A, B). As rupture events can no longer be visualized after all beads are ejected from a sphere, we injected 5 h post amputation to track rupture events during LPOs or 24 h post amputation to track ruptures during SPOs. We observed that rupture sites are randomly distributed in spheres injected at 5 h (Fig. 2C i), but significantly more localized in spheres injected at 24 h (Fig. 2C ii). We compared rupture site locations for both LPOs and SPOs to data drawn from a uniform distribution using a two sample Kolmogorov-Smirnov test, and found that the 5 h data are not significantly different from a uniform distribution (p = 0.9702) while the 24 h data are (p = 1.0047e-07.) This suggests that a mechanical weak spot in the tissue sphere forms as regeneration proceeds.

**Figure 2.**
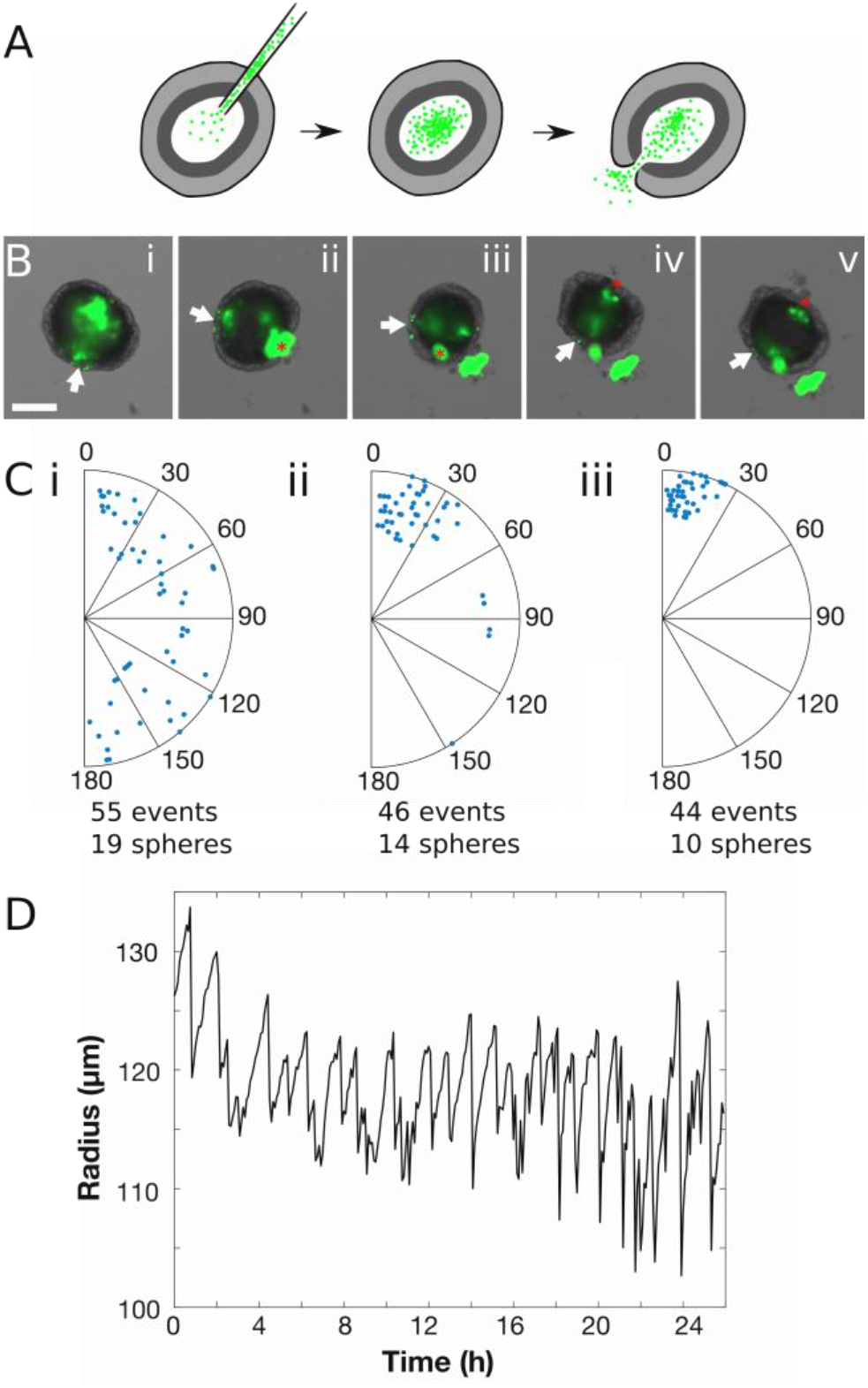
Rupture site becomes constant with head development. A. Experiment schematic showing injection of fluorescent microbeads into a hollow sphere. Beads are ejected from the sphere during rupture events. B. Representative image series of sphere ejecting beads during successive ruptures. White arrow indicates the feature used to track rotation, red asterisk represents observed rupture site (see Materials and Methods). Scale bar 100 µm. C. Location of the rupture site relative to the first rupture, each radius representing a single sphere. i. Beginning 5 h after cutting. ii. Beginning 24 h after cutting. iii. Head pieces, containing the mouth of the parent animal. D. Representative oscillation plot of a head tissue piece.

To confirm that this structural weak point corresponded to the *Hydra* mouth, we tracked ruptures in regenerating head tissue pieces containing the intact mouth of the parent animal. We found that these head piece spheres had an invariant rupture site (Fig. 2C iii), supporting the idea that the emergence of a fixed rupture site is coincident with mouth development during regeneration. Finally, to confirm a link between the mouth and oscillation dynamics we analyzed the oscillations of head pieces and found that they only exhibit SPOs as seen from the distribution of time periods (Fig. 2D, Table 1). These data demonstrate that the presence of a mouth in a tissue piece is sufficient for SPOs.

As it had been proposed that the aboral pore acts as a second weak point in the intact animal that may be used for pressure regulation (35), we also imaged foot tissue pieces containing the entire basal disc (Fig. 1A). Foot tissue pieces showed oscillation parameters with a greater similarity to LPOs than to SPOs (Table 1). The statistically significant difference in period between foot tissue pieces and SPOs indicates that the presence of an aboral pore does not increase rupture frequency in the same way the presence of a mouth does. Thus, the aboral pore does not play a role in regulating osmotic pressure during regeneration. We suspect the similarity of swelling rate to that of SPOs results from a difference in tissue composition in the foot. Both the hypostomal region and the basal disc have significantly higher proportions of epitheliomuscular and nerve cells than the body column (36), which may cause differences in mechanical properties or permeability.

In summary, these results support the hypothesis that ruptures during LPOs are caused by osmotically-driven inflation until the yield strength of the tissue is reached, resulting in random rupture locations. In contrast, SPOs are due to the development of a mouth structure, creating a permanent, localized weak point on the sphere. This is consistent with previous observations that insertion of head tissue into cell aggregates decreases the time required for a shift to SPOs to occur (18). The presence of a head organizer would allow the aggregate to more rapidly define a head and develop a mouth, resulting in a faster oscillation pattern shift. Whether the forming mouth acts solely as a mechanical defect as previously suggested (14), or actively regulates osmotic pressure cannot be distinguished based on these data.

### Mouth function is required for a shift to SPO

To determine whether the mouth plays an active role in osmoregulation in regenerating tissue spheres, we decouple mouth function from mouth structure by examining nerve-free *Hydra*, which are capable of complete regeneration but are unable to open their mouths to relieve pressure or respond to chemical stimuli (31, 37, 38). In contrast to normal animals, nerve-free *Hydra* take on a characteristic bloated appearance (22)(Fig. 3A) due to their inability to relieve internal pressure by mouth opening. The mouth appears morphologically normal in nerve-free animals (Fig. 3B), suggesting that lack of function is caused by the absence of neurons and thus an inability to sense pressure (39). Body column tissue spheres derived from nerve-free animals showed only LPOs, with a period slightly longer than LPOs in wild-type spheres (Fig 3C, Table 2). The small difference in parameters may be due to differences in tissue strength given that nerve-free animals lack all cell types derived from the interstitial cell lineage: neurons, gland cells and nematocytes (40). The complete lack of SPOs in tissue pieces derived from body columns of nerve-free *Hydra* implies that either nervous system control of mouth opening is required to transition to SPO or development of a mouth structure is delayed in nerve-free *Hydra*. Nerve-free animals regenerate, suggesting that nerve cells are not critical (41). However, regeneration defects occur in wild-type animals when neurogenesis is inhibited (42), suggesting that nerve-free animals may use an alternate regeneration pathway for regeneration. Since we only record oscillations for 48 h, it is possible that we would not observe the pattern shift if its occurrence was significantly delayed in nerve-free animals compared to the wild-type.

**Table 2.**
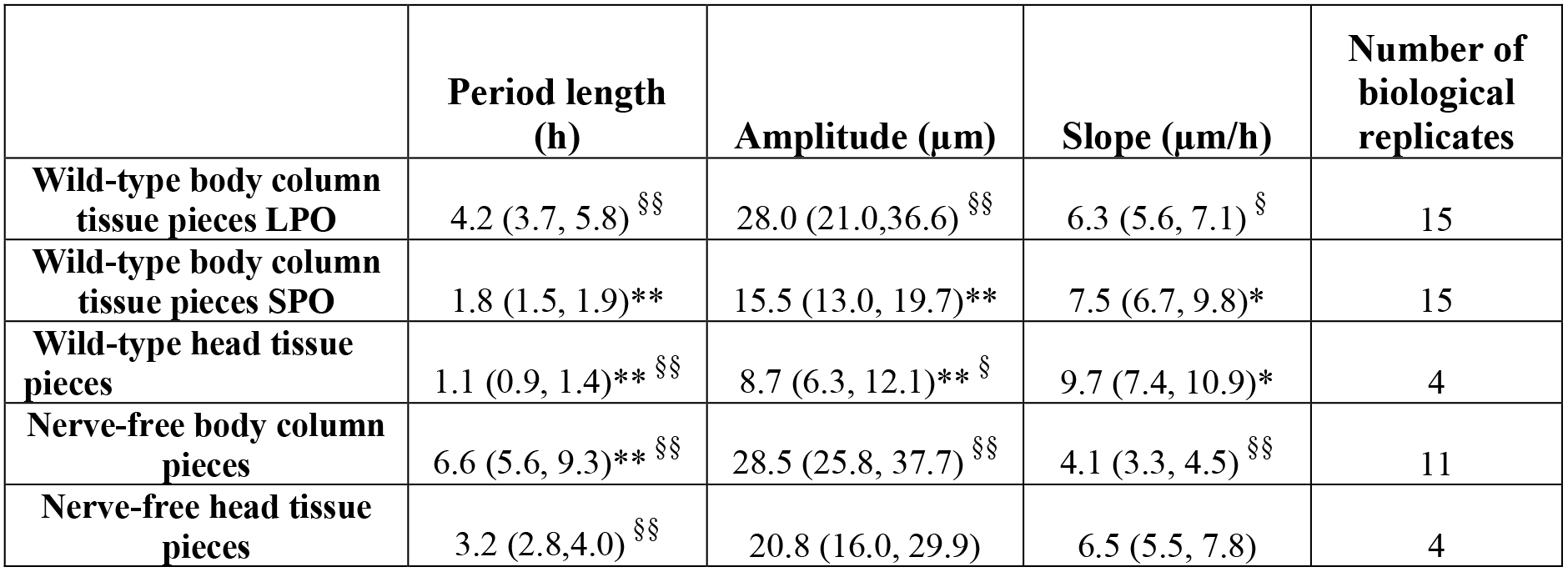
Oscillation parameters for various experimental conditions. Parameters are reported as median of biological replicates with the first and third quartiles. * indicates significant difference from wild-type LPOs at p < 0.05. ** indicates significant difference from wild-type LPOs at p < 0.01. ^§^ indicates significant difference from wild-type SPOs at p < 0.05. ^§§^ indicates significant difference from wild-type SPOs at p < 0.01.

**Figure 3.**
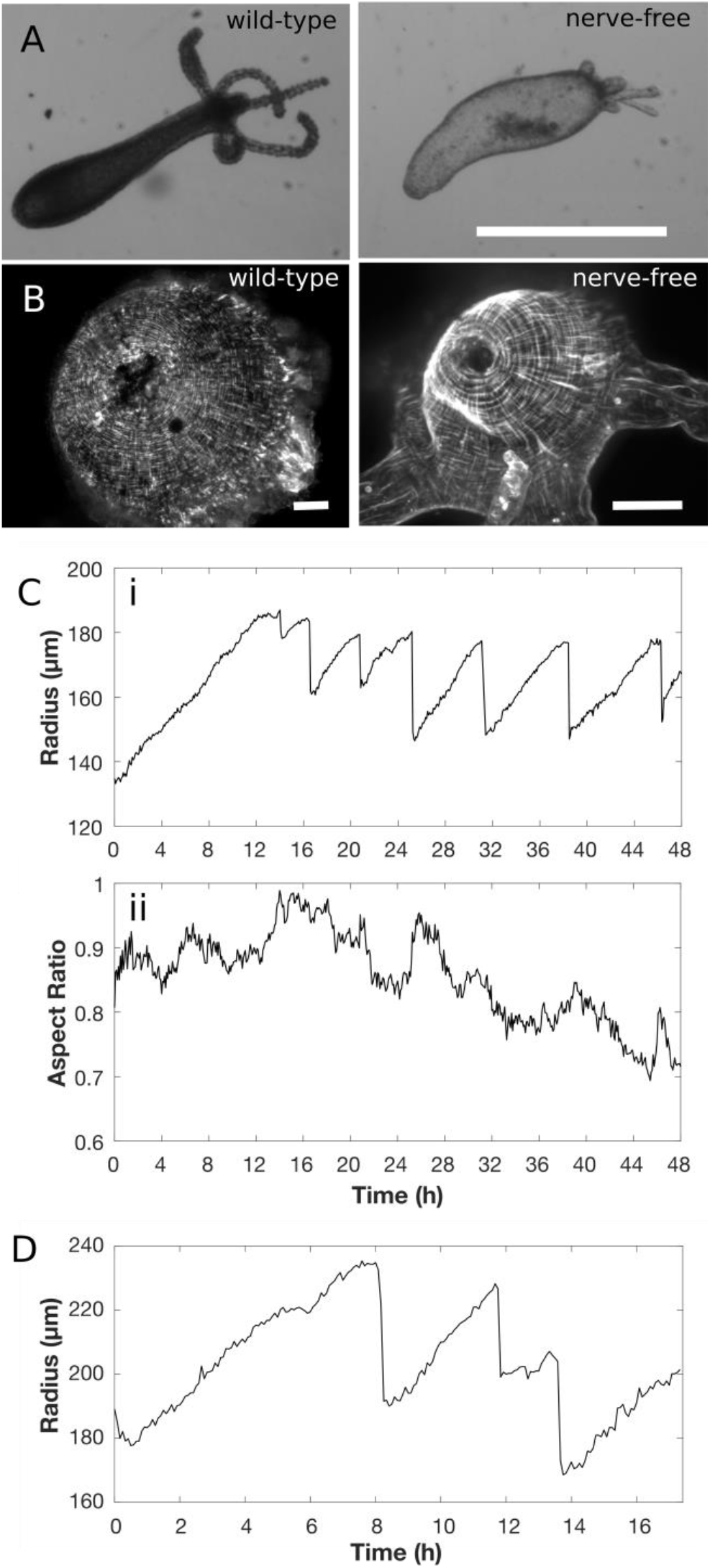
Mouth opening impacts oscillation dynamics. A. Comparison between wild-type and nerve-free polyps, showing characteristic bloated phenotype of nerve-free *Hydra* (scale bar 1 mm). B. Comparison of myoneme organization in the hypostome of wild-type and nerve-free animals (scale bars 50µm). C. Representative plots of a nerve-free tissue sphere. i. oscillation plot, ii. aspect ratio. D. Representative oscillation plot of nerve-free head tissue sphere.

Therefore, interpreting the unusual oscillation pattern in nerve-free spheres requires us to fully decouple mouth structure from mouth function. We achieve this using tissue pieces cut from the heads of nerve-free animals. As these contain a fully-formed mouth, they bypass the possibility of delayed head regeneration in nerve-free *Hydra*. If the presence of a mouth structure was sufficient to increase rupture frequency, we would expect to observe SPOs in spheres derived from nerve-free head pieces as we do in untreated wild-type head pieces (Fig. 2D). Instead, we find that spheres from nerve-free head pieces show only LPOs (Fig. 3D, Table 2).

Taken together, these data demonstrate that the shift in oscillation pattern observed in regenerating *Hydra* tissue spheres is caused by the formation of a functional mouth and its use in osmoregulation. A tissue sphere derived from a wild-type polyp initially exhibits LPOs, where rupture is dictated by yield strength of the tissue. Rupture events in this regime are random as mechanical failure is equally likely to occur at any point on the sphere. Approximately 24 hours into the regeneration process, we observed the development of a functional mouth which allows for active osmoregulation, causing the shift to SPOs.

### Implications for theoretical models of *Hydra* regeneration

Various attempts have been made to model axis determination from a homogenous initial state in *Hydra* spheres. The core of these models lies in some form of feedback between morphogen concentrations and mechanical properties of the tissue such as elasticity. The dynamics of the morphogen concentrations are modelled using the Gierer-Meinhardt model (3), while feedback between mechanics and the morphogens is modelled using a relation between tissue stretch and morphogen diffusion. In the model proposed by Soriano *et al.*, this takes the form of a linear relationship between tissue strain and the diffusion coefficient of one of the morphogens. Axis formation is posited to occur when a stable gradient is established - a consequence of the diffusion coefficient exceeding a certain threshold (18), which makes the timescale for gradient formation much shorter than the timescale for shape oscillations. In a more recent model by Mercker *et al.* the local diffusion coefficient is a function of the local area strain of the tissue and the elastic modulus is a function of morphogen concentration. This allows for a growth instability - high local strain causes accumulation of the morphogen and morphogen accumulation allows for higher local strains in response to the same stress (4).

To date there are no quantitative experimental data on concentration patterns of morphogens, their diffusion constants, or the feedback between morphogen concentration and mechanical properties in *Hydra*. Therefore, models rely entirely on relations between morphological parameters such as swelling rate, initial tissue size, and the time of shape symmetry breaking to constrain model parameters and validate predictions. Some of these constraints can be independently verified using our data. For example, the model proposed by Mercker *et al.* assumes that the tissue sphere deflates instantaneously to its initial volume if the local strain exceeds a certain threshold value and then heals without a change in the volume over a certain period of time. From our oscillation parameters, we can calculate the area strain threshold during LPOs in body column tissue spheres using the ratio of amplitude to minimum radius. We estimate that it is approximately (1.23)^2^, which is close to the value of (1.25)^2^ used by Mercker *et al*.(4).

In contrast, other model constraints are inaccurate in light of our data. For example, the results presented here force us to reconsider the assumption that the time of symmetry breaking always coincides with the time of the oscillation pattern shift from LPO to SPO for two reasons: First, spheres derived from body column tissue pieces show the pattern shift despite inheriting the parental body axis (19). Second, nerve-free tissue pieces only exhibit LPOs but nevertheless break symmetry (Fig. 3C). Thus, the time of oscillation pattern shift is not necessarily the time of symmetry breaking, and one of the key observables used to constrain the models is not universally applicable. Instead, we show that the shift in the oscillation pattern is caused by a change in local yield strength of the tissue due to mouth formation, a property whose variation existing models do not consider.

We estimate the local yield strength of the tissue by treating it as an elastic shell (see Materials and Methods). The order of magnitude estimate is made using only quantities that can be measured or calculated from experimental data presented here or elsewhere in the literature, except for Poisson’s ratio which does not affect the order of magnitude (see Materials and Methods). The estimated elastic pressure inside the sphere at the time of rupture is on the order of 10 Pa during LPOs. Since the pressure scales linearly with oscillation amplitude (see Materials and Methods) and the SPO amplitude is approximately half the LPO amplitude, the pressure at the time of rupture during SPOs is approximately 5 Pa acting on an area of the order of 2-3 cell diameters across. Therefore, the elastic force must be on the order of a few nanoNewtons at the time of rupture during SPOs. The magnitude of this force is comparable to that exerted by myonemes to create a mouth opening (33) and to the separation force associated with tight junctions involved in cell-cell adhesion (43). While the sources of the elastic forces estimated here for SPOs are different from those involved in mouth opening, they act on the same tissue producing the same effect – breaking cell-cell contacts to create an opening, suggesting that the estimates are reasonable. We thus provide an experimentally determined value that can be used to constrain parameters associated with tissue rupture in models.

Current models assume that mechanical forces on the *Hydra* sphere during oscillations drive morphogen patterning, predicated largely on the assertion that oscillation pattern shift and axis specification are closely linked. The most compelling evidence in favor of this theory is the observation that completely suppressing oscillations via increased medium osmolarity prevents regeneration (18), though this cannot distinguish between lack of oscillations and osmotic stress on a freshwater animal as the cause. Furthermore, mechanotransduction pathways are known to be involved in a wide range of morphogenetic and developmental processes (44). β-catenin, which acts as a mechanotransducer in other model organisms (45), is involved in *Hydra* head specification via the canonical Wnt pathway (46, 47). Consequently, it has previously been proposed that β-catenin may link the mechanical forces due to tissue stretch or rupture with biochemical patterning in *Hydra* (18). This remains to be experimentally verified. The timing of mRNA expression of various components of the canonical Wnt pathway during *Hydra* regeneration after injury or from spheres has been investigated by *in situ* hybridization (14, 15). However, this only evaluates transcription on fixed animals and thus these data may not be accurate representations of protein localization and *in vivo* dynamics of regeneration. Our results suffer from a similar limitation: we have shown that the oscillation pattern shift is a consequence of mouth function and thus that axis specification occurs earlier than previously assumed but cannot directly measure the underlying morphogenetic processes.

Developing and testing a model that fully describes pattern formation in regenerating *Hydra* spheres requires quantitative data of the sort that morphological readouts alone cannot provide. The advent of modern tools and techniques offers the ability to observe and interact with the system in novel ways. Future studies should aim to combine pharmacological perturbations to probe protein interactions with mechanical perturbations to alter the forces during oscillations, and monitor the effects of these perturbations on axis specification via simultaneous use of transgenic reporter strains to visualize morphogen gradients *in vivo.* Quantification of morphogen concentrations and dynamics is necessary for a full understanding of the biochemistry in a way that is currently impossible. Finally, while this and other recent works (10, 13, 19) have focused on regenerating spheres originating from tissue pieces, the oscillation behavior of spheres originating from cell aggregates should be revisited. A direct comparison of the results from these two starting scenarios is likely to provide further insights into the mechanisms that drive regeneration and patterning. Exploring these possibilities and leveraging them to improve existing models should be the next step in our attempt to understand axis specification in *Hydra*.

## CONCLUSIONS

During *Hydra* regeneration from small tissue pieces or aggregated cells, a hollow bilayered sphere forms that undergoes dramatic shape oscillations. A switch in oscillation pattern, from long period, large amplitude to short period, small amplitude oscillations occurs approximately one day into regeneration. As previous explanations for the shift in oscillation pattern have recently been invalidated, we reexamined this fundamental process during *Hydra* regeneration from tissue spheres and demonstrate that the oscillation pattern shift is a direct consequence of the onset of mouth function and its use in osmoregulation. This allows us to infer the development of an important physiological function through a morphological read out. The results from this work also enable the field to reexamine and improve existing models of *Hydra* regeneration that rely on the concurrence of the shift in oscillation pattern and decrease in aspect ratio to constrain model parameters.

## FUNDING INFORMATION

This work was funded by NSF grant CMMI-1463572 (E.-M.S.C.), the Research Corporation for Science Advancement (E.-M.S.C.), the Gordon and Betty Moore foundation (E.-M.S.C.), and US Department of Energy Grant FG02-04ER54738 (P.H.D.).

## AUTHOR CONTRIBUTIONS

E.-M.S.C. designed research. R.W. and T.G. performed experiments. R.W., T.G., K.K., H.J.Q. and Z.S. analyzed data. K.K. contributed analytical tools. P.H.D. consulted on data analysis and interpretation. R.W., T.G., P.H.D., and E.-M.S.C. wrote the manuscript.

## Acknowledgements

The authors thank Cassidy Tran for providing nerve-free *Hydra*, Elizabeth Lanphear, Christina Rabeler, Sara Martin and Connor Keane for help with *Hydra* care, Winnie Shi and Haochen Wang for help with the data analysis, Dr. Olivier Cochet-Escartin for discussion, and Dr. Rob Steele for discussion and comments on the manuscript. We also thank Dr. Hiroshi Shimizu for providing us the sf-1 strain, and Dr. Alison Hanson for the A10 strain.

